# Dynamic DNA methylation contributes to carryover effects and beneficial acclimatization in geoduck clams

**DOI:** 10.1101/2022.06.24.497506

**Authors:** Hollie M. Putnam, Shelly A. Trigg, Samuel J. White, Laura H. Spencer, Brent Vadopalas, Aparna Natarajan, Jonathan Hetzel, Erich Jaeger, Jonathan Soohoo, Cristian Gallardo-Escárate, Frederick W. Goetz, Steven B. Roberts

## Abstract

Given the rapidly changing global climate, there is a growing need to understand the acclimatory basis of animal response to environmental change. To investigate the role of DNA methylation in environmental acclimatization, we generated a reference genome and surveyed the physiology and DNA methylomes of juvenile geoduck clams, *Panopea generosa,* under differing seawater pH regimes. Juveniles were initially exposed to one of three seawater pH conditions, followed by ambient common-garden conditions, then a second reciprocal exposure to ambient pH or pH 7.4. Within 10 days of the initial low pH exposure, juvenile clams showed decreased shell size relative to ambient pH with corresponding differential DNA methylation. Following four months of ambient common-garden conditions, juveniles initially exposed to low pH compensatorily grew larger, with DNA methylation indicative of these phenotypic differences, demonstrating epigenetic carryover effects persisted months after initial exposure. Functional enrichment analysis of differentially methylated genes revealed regulation of signal transduction through widespread changes in the Wnt signaling pathways that influence cell growth, proliferation, tissue and skeletal formation, and cytoskeletal change. After 10 days of secondary exposure to pH 7.4, naive juvenile clams were more sensitive to low pH compared to those initially exposed, showing reduced growth and having nearly a 2-fold greater change in DNA methylation. Collectively, this new genomic resource and coordinated phenotypic and methylomic response support that epigenetic mechanisms underlie acclimatization to provide beneficial phenotypes.

## INTRODUCTION

### Mollusc genomics to inform acclimatization and adaptation

Of metazoans, Mollusca is the second most species-rich phylum and despite their critical economic, ecological, and scientific importance genomic resources available for Mollusca lag far behind other metazoan lineages (Gomes-dos-Santos et al. 2020). High-quality genomic resources can help address questions about diversity, evolution, phylogeography, demography, adaptation patterns, and conservation traits (Savolainen et al. 2013; Dunn and Ryan 2015; Richards 2015; Lopez et al. 2019; Arumugam et al. 2019), all of which would contribute to preserving the economic and ecological services molluscs provide. With large population sizes, wide habitat distribution, and high fecundity rates, the high heterozygosity and repetitive sequence context of molluscan genomes has been a long-standing challenge in genome assembly, though there currently are a number of genomes available including those of the Pacific oyster, *Crassostrea gigas* (Zhang et al. 2012), the Eastern oyster, *Crassostrea virginica (Gómez-Chiarri et al. 2015)* and the Manilla clam, *Ruditapes philippinarum* (Yan et al. 2019). Challenges in genome assembly have been overcome by the combination of short-read sequencing and long read, or linked-read sequencing (Li et al. 2018a; Zhang et al. 2012), which has enabled the generation of long scaffolds corrected by extensive short reads. Existing marine bivalve genomic resources have been used to improve breeding programs and aquaculture productivity (Nguyen et al. 2014; Kijas et al. 2018; Masonbrink et al. 2019). Further, these resources have enabled investigations of stress adaptation mechanisms that lead to identification of the underpinnings of environmental tolerance and disease resistance (Murgarella et al. 2016; Mun et al. 2017; Li et al. 2018a; Powell et al. 2018). With ongoing and impending climate change, including the negative impacts of ocean acidification already observed in marine bivalves (Yuan et al. 2020; Fitzer et al. 2018; Ramesh et al. 2017; Speights et al. 2017; Waldbusser et al. 2011; Gazeau et al. 2007; Stumpp et al. 2011; Pan et al. 2015; Boulais et al. 2017; Byrne and Przeslawski 2013), and consequent negative impacts on marine ecosystems at large (Marshall et al. 2017; Busch and McElhany 2016; zu Ermgassen et al. 2013), it is becoming more critical these high-quality genomic resources be developed and experiments be carried out to understand mechanisms of environmental tolerance.

### Geoduck and environmental tolerance through priming

The Geoduck clam, *Panopea sp.,* is an ideal species for studying environmental tolerance because in addition to being the basis of a high-value commercial fishery, it is among the longest lived animals on earth despite being a sessile marine invertebrate that needs to acclimate and adapt to dynamic local conditions. While the Pacific geoduck, *Panopea generosa*, has shown delayed development under low pH stress (Timmins-Schiffman et al. 2020), interestingly it has also shown compensatory growth when re-exposed to low pH stress to which it was previously conditioned (Gurr et al. 2020). This beneficial acclimation suggests conditioning or ‘priming’ could enhance environmental tolerance and there is a growing body of evidence supporting this concept in a variety of biomineralizaing marine invertebrate species (Eirin-Lopez and Putnam 2019), including the Sydney rock oyster (Parker et al. 2015), the Manilla clam (Zhao et al. 2018), and the purple sea urchin (Strader et al. 2019).

### Methylation as a mechanism of environmental tolerance and priming

One proposed molecular mechanism of environmental tolerance is environmentally-induced DNA methylation, which can regulate gene expression (Mirbahai and Chipman 2014). DNA methylation is the addition of a methyl group on a cytosine nucleotide typically found in a CpG context. In marine invertebrates, gene body DNA methylation has a positive relationship with gene expression and a negative relationship with gene expression variability (Li et al. 2018b; Liew et al. 2018; Gavery and Roberts 2013), indicating that environmentally-induced DNA methylation changes impact gene expression and therefore phenotype. Increasing efficiency of gene expression regulation through DNA methylation may be a particularly important mechanism to reduce energy demands in developing and sessile marine invertebrates. The gold standard of DNA methylation analysis includes bisulfite conversion of DNA followed by sequencing, to generate single nucleotide level resolution for hypothesis testing around the interaction of genome features and DNA methylation. Studies have found DNA methylation changes in gene body regions in Eastern oysters exposed to low pH conditions through bisulfite sequencing approaches, suggesting a role of DNA methylation in gene regulation and acclimatization (Venkataraman et al. 2020; Downey-Wall et al. 2020). There is however, a need to understand mechanisms of stress priming at the molecular level (Hackerott et al. 2021; Costantini 2014; Costantini et al. 2010). This knowledge could lead to more effective progress in resilience husbandry for food production and conservation restoration systems. To this end, we: 1) sequenced, assembled, and annotated a draft genome for the Pacific geoduck clam, *Panopea generosa*, and 2) designed a multi-stage reciprocal exposure experiment to test the hypotheses that initial exposure to a low pH would induce differential DNA methylation and priming of epigenetic “memory”, resulting in an acclimatized phenotype.

## RESULTS

### A Panopea generosa genome

The Proximo Hi-C assembly process (Phase Genomics) resulted in a set of 18 chromosome-scale scaffolds containing 1.42 Gbp of sequence (64.53% of the corrected assembly). Juicebox correction resulted in a final set of 18 scaffolds spanning 942 Mbp, with a scaffold N50 of 57,743,597 bp and a scaffold N90 of 34,923,512 bp (**Figure 1** and **Table 1**). Genome annotation identified 34,947 genes and 236,960 coding sequence (CDS) regions, corresponding to 38,326 mRNA features (**Table 1**). Genome feature tracks including those characterizing genes, exons, introns, repetitive sequences (including predicted transposable elements), and CG motifs are available (Roberts et al. 2020). The assembled genome is available on NCBI under GCA_902825435.1.

**Figure 1.**
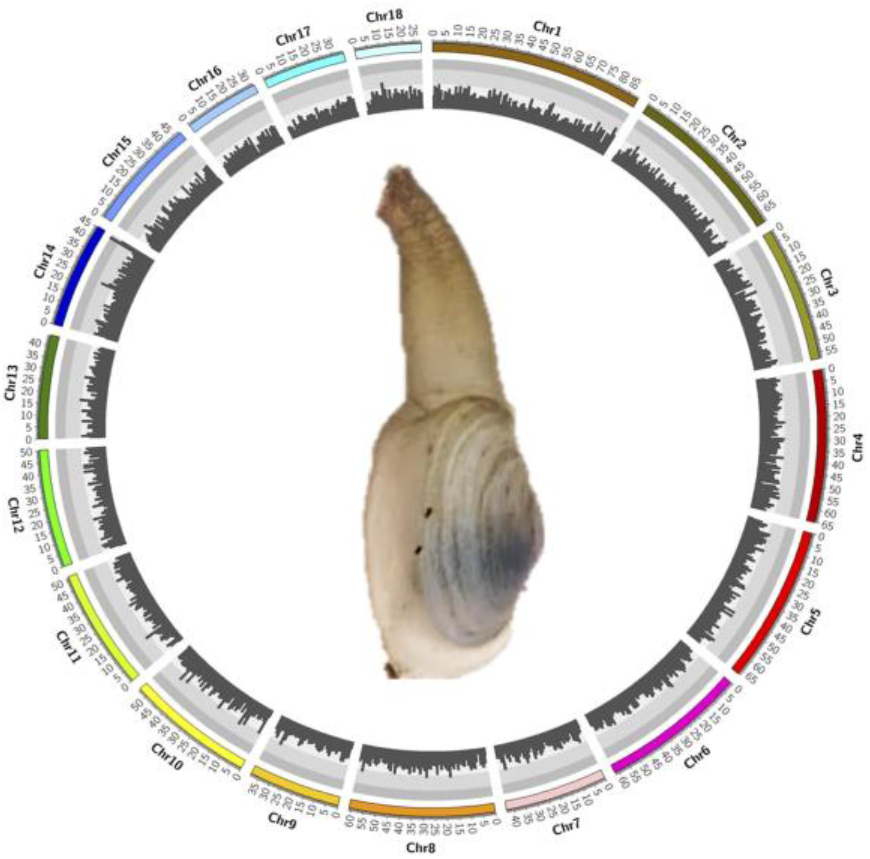
Genome circos plot showing 18 chromosomes with chromosome name on the outermost track, followed by chromosome position in kilobase pairs (kbp), color bars indicating each chromosome, and gene density (genes/kbp) on the innermost track.

**Table 1.** Summary table of assembled and annotated Panopea generosa genome features including their counts and size in base pairs (bp). Unidirectional arrangement of genes refers to the frequency that genes on either side of an intergenic region are in the same direction.

|  |  |  |
| --- | --- | --- |
|  | Total scaffold length (bp) | 942,353,201 |
| **Genome** | Scaffold N50 (bp) | 57,743,597 |
|  | GC content, N excluded (%) | 33.78 |
|  | Number of genes | 34,947 |
| **Genes** | Mean gene length (bp) | 201 |
|  | Mean coding region length (bp) | 201 |
| **Exons** | Mean number of exons per gene | 6.78 |
|  | Mean length (bp) | 201 |
|  | Total length (Mbp) | 47 |
|  | Genes with introns (%) | 72.04 |
| **Introns** | Mean length (bp) | 2185 |
|  | Total length (Mbp) | 338 |
|  | Predominant first two nucleotides at donor splice sites | TA (48%)<br>TG (42%) |
| **Intergenic** | Mean length (bp) | 16,394 |
|  | Unidirectional arrangement of genes | 49.5% |

Genome analysis with BUSCO against the Metazoan dataset identified 71.5% complete, single-copy orthologs (Complete: 71.5% [Single copy: 64.1%, Duplicated: 7.4%], Fragmented: 8.3%, Missing: 20.2%, n=978 BUSCO reference genes). Genes had a mean length 10,811bp and median length of 4,464bp. The 38,326 mRNAs identified had a mean length of 12,903bp and median length of 5,453bp. Annotation yielded 16,889 tRNAs with a mean and median length of 75bp with the shortest being 53bp, and the longest 314bp. A total of eight rRNA were identified with a mean length of 117bp, median length of 115bp and a range of 113 - 138bp. Repetitive sequence analysis indicated that 40.24% of the assembled genome contains repeats with a total of 1,676,544 transposable elements (**Table S1**). GC content was determined to be 33.78% and a total of 15,712,294 CG motifs are present in the genome.

### Comparative genomics

In comparing the *Panopea generosa* genome to the genomes of other bivalve species, we found that clam genomes tend to be ~1.5-2 times larger in size than oyster genomes, consisting of almost twice as many putative chromosomes (**Table 2**). Despite this difference in size and number of putative chromosomes, the number of genes tended to be similar across all species.

**Table 2.** Comparison of bivalve genome features.

| Species | GenBank Assembly Accession | Genome size (Mbp) | Number of putative chromosomes | Number of Genes | GC content (%) | Reference |
| --- | --- | --- | --- | --- | --- | --- |
| *P. generosa* | GCA_902825435.1 | 942 | 18 | 34,947 | 33.78 | This Study |
| *R. philippinarum* | GCA_009026015.1 | 1230 | 19 | 27,652 | 32.26 | (Mun et al. 2017) |
| *C. virginica* | GCA_002022765.4 | 685 | 11 | 38,826 | 34.82 | (Gómez-Chiarri et al. 2015) |
| *C. gigas* | GCA_902806645.1 | 648 | 10 | 38,296 | 33.50 | (Zhang et al. 2012) |

### Experimental design and data collection

To examine growth phenotype and DNA methylation patterns across time and environments varying in pH, we initially exposed juvenile geoduck to three different pH conditions (7.9 [ambient], 7.4, and 7.0) for 23 days. To examine the persistence of any effects on phenotype and methylation from the initial treatment, clams were next held under ambient common garden conditions, while keeping initial treatment groups separated, for 112 days (**Figure 2**). Finally, to investigate any effects of the initial treatment on the phenotypic and methylation response to low pH re-exposure, clams from each initial treatment group were re-exposed in a reciprocal fashion to pH 7.9 or 7.4 for an additional 23 days. Samples were collected for size and molecular analyses.

**Figure 2.**
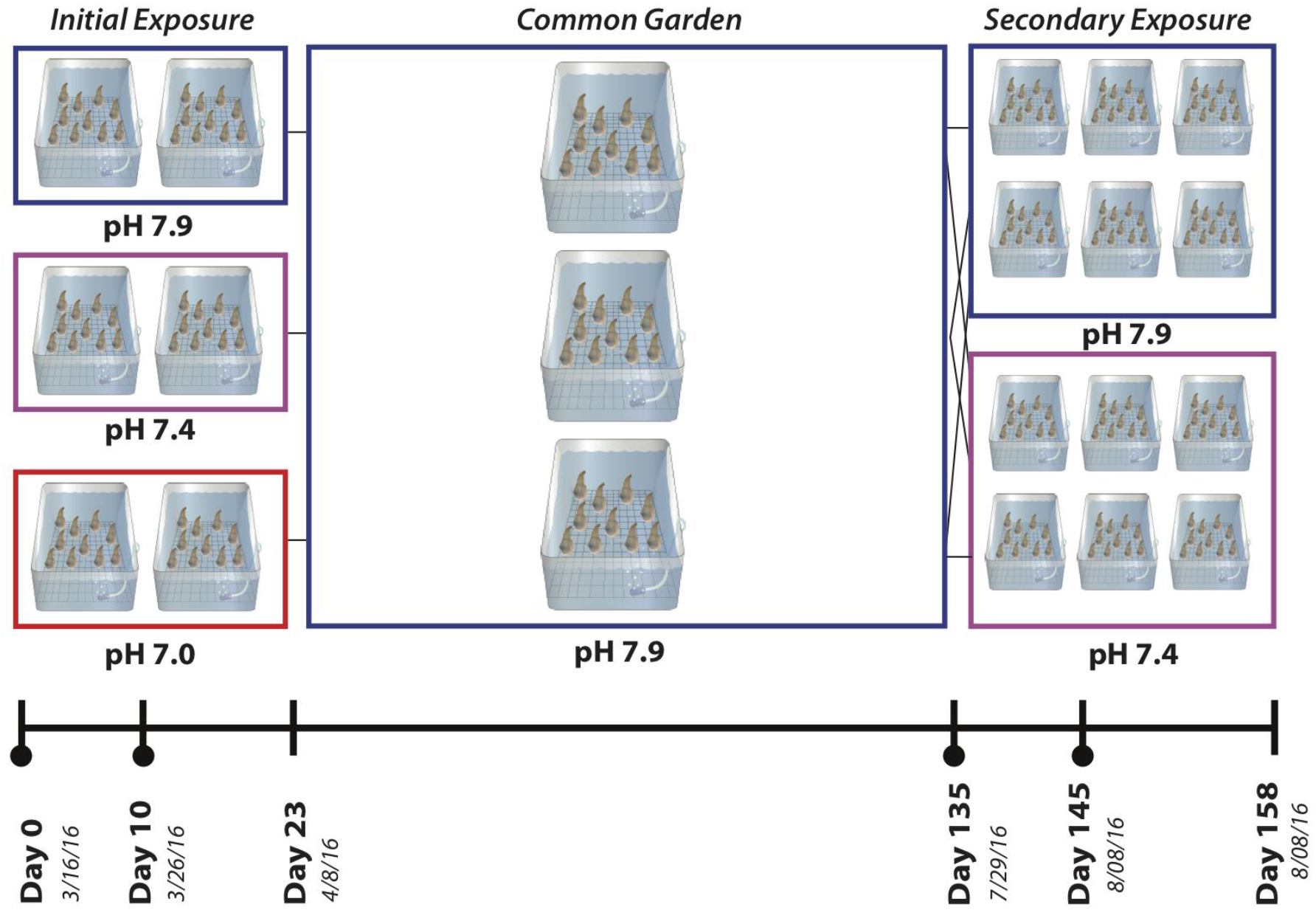
Experimental design and sampling time point for geoduck low pH exposure. Phenotypic sampling was performed on days indicated by the vertical lines on the timeline, whereas molecular sampling was performed on days indicated by the black circles below the vertical lines.

### Low pH exposures show different effects on growth

In general, clams increased in shell size (shell area), regardless of rearing condition, over the first 145 days of the experiment. Survivorship was high across all pH conditions, with no mortality during different pH exposures. There was no significant interaction between initial pH exposure and time (p>0.05) on relative shell size. However, there was a significant effect of pH on relative shell size (F_2,96_=12.436, p<0.00001) after 10 and 23 days of initial exposure, with a 21.4% and 29.2% lower mean relative shell size, respectively, at pH 7.0 compared to pH 7.9 (**Figure 3A**). There was a significant effect of time (F_2,96_=18.202, p<0.00001) on relative shell size, with relative shell size increasing regardless of pH exposure within the 23 days (**Figure 3A**). After the common garden (Day 135), there was again no significant interaction effect between pH and time on relative shell size, and all clams showed an increase in size over this period. However, clams previously exposed to low pH treatments had significantly larger mean relative shell sizes than those previously exposed to ambient conditions (F_2,132_=23.113, p<0.00001), with clams initially exposed to low pH (pH 7.4 and 7.0) having 1.4 and 1.3 times larger relative mean shell sizes, respectively, than those initially exposed to ambient pH (pH 7.9) (**Figure 3B**). After both 10 and 23 days of secondary pH exposure (Day 145 and Day 158, respectively), there was a significant interaction effect between initial and secondary pH exposure (F_2,123_=5.339, p=0.0059) (**Figure 3C and D**). This interaction effect was driven by a more severe negative growth effect in naive clams exposed to pH 7.4, which showed smaller mean relative shell size than clams initially exposed to low pH (pH 7.4, 7.0) (**Table S2**).

**Figure 3.**
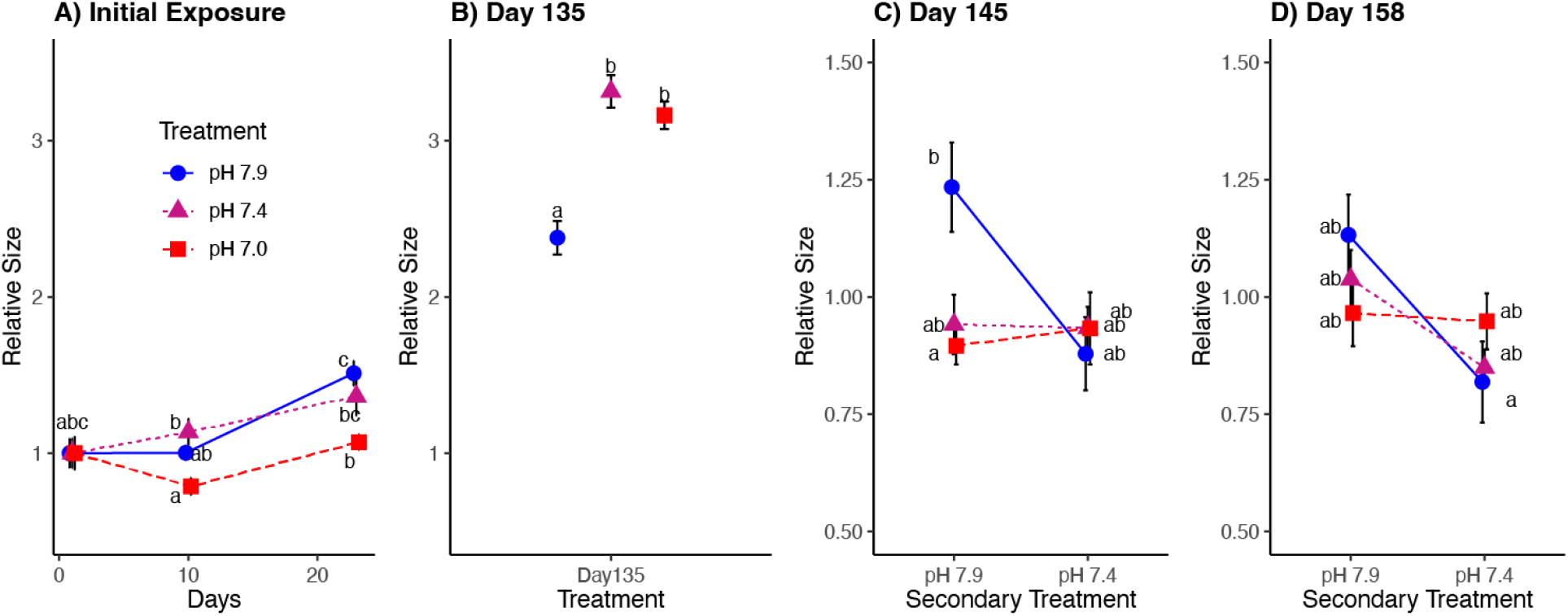
Relative shell size (mean ± sem) after (A) the initial exposure, (B) the ambient common garden period (Day 135), (C) 10 days into the secondary reciprocal exposure (Day 145), and D) 23 days into the secondary exposure (Day 158). Results of posthoc analyses are indicated by dissimilar letters for significant pH effects.

### Methylation Results

A total of 52 samples were sequenced and raw data are available in the NCBI Sequence Read Archive (SRA; BioProject: PRJNA566166; Accessions: SRR10143613 - SRR10143660). These libraries generated 1.2 billion sequence reads with a mean of 23.8 ± 5.5 million reads per sample. The mean bisulfite conversion efficiency across all samples was 98.83%, as calculated from the lambda phage spike-in. Summary tables for trimming and alignment statistics can be found in **Tables S3** and **S4**, respectively.

In order to assess the influence of altered pH conditions on DNA methylation in geoduck clams, two complementary strategies were used - differentially methylated region (DMR) and differentially methylated gene (DMG) analysis. While a growing body of work has been focused on gene level analyses of methylation in invertebrates, this does not preclude significant changes of methylation in other regions that would be missed by a gene-centric approach. After 10 days of initial pH exposure, there were 41 DMRs ranging from 21 to 1701 base pairs in length (**Figure S1A**, **Table S5**). Of these 41 DMRs, 8 (19.5%) showed at least a 2-fold decrease in mean methylation and 12 (29.3%) showed at least a 2-fold increase in mean methylation at pH 7.4 relative to ambient conditions (pH 7.9) (**Figure S2A**). Contrastingly at pH 7.0, a larger proportion of DMRs showed a decrease in mean methylation (9, 22.0%) and only 6 (14.6%) DMRs showed an increase in mean methylation relative to ambient (**Figure S2A**). Following common garden rearing at Day 135, we found a total of 22 DMRs ranging from 28 to 337 base pairs that generally showed similar trends in methylation changes across initial low pH treatments relative to ambient, which was predominantly a decrease in methylation (**Figure S1A**, **Figure S2B**, and **Table S6**). Of the 1627 and 740 regions analyzed 10 days after the initial pH exposure (Day 10) and after the common garden (Day135), respectively, there were 524 regions that overlapped although none showed significantly different methylation (*P* < 0.05 by one-way ANOVA) with respect to pH at both time points. Following 10 days of secondary low pH exposure (Day 145), we found a total of 42 DMRs ranging from 47 to 891 base pairs, with naive clams mostly showing an increase in methylation upon low pH exposure and ‘primed’ clams showing subtler methylation changes (**Figure S1A**, **Figure S2C,** and **Table S7**). DMRs from all comparisons tended to be broadly distributed across the genome and along scaffolds, with no apparent spatial trend (**Figure S1B-C**). DMRs from after 10 days of initial pH exposure occurred significantly more in putative three-prime untranslated regions, where DMRs at Day 145 occurred significantly more than expected in introns (**Figure S3**, **Table S8**). Overall, DMR analyses showed pH affected methylation particularly around and within genes.

### Differentially Methylated Genes (DMGs)

Given the demonstrated gene body methylation in bivalves (Roberts and Gavery 2012) and other marine invertebrates (Sarda et al. 2012) we tested for differentially methylated genes (DMGs) across pH exposures at different sampling points.

Following 10 days of exposure to low pH, 1,417 genes were differentially methylated between pH treatments (**Table S9**). GO enrichment analysis identified 102 enriched terms (**Table S10**) with regulation of cell morphogenesis showing the most significant over representation (p=0.0017) along with another 5 GO terms under the GOSlim category of cell organization and biosynthesis (**Table S10**). Additional enriched GO terms were related to the Biological Process of cell cycle and proliferation, developmental process, DNA metabolism, protein metabolism, signal transduction, transport, stress response, other biological processes, other metabolic processes.

Following the ambient common garden at Day 135, there were 797 DMGs due to initial pH exposure (**Table S11**). GO term enrichment identified that oxidative DNA demethylase activity and DNA demethylation were the most significantly overrepresented terms at p=0.0016 and p=0.005, respectively. Other enriched GO terms were related to the Biological Processes cell cycle and proliferation, cell organization and biogenesis, cell-cell signaling, death, developmental processes, DNA metabolism, protein metabolism, RNA metabolism, signal transduction, stress response, transport, other biological processes, and other metabolic processes (**Table S12**).

Following secondary pH exposure at Day 145, there were 434 DMGs showing an interactive effect of initial and secondary treatment (**Table S13**). These DMGs showed a pattern similar to shell area, where naive clams exposed to the low pH (initial pH 7.9, secondary pH 7.4) showed a 2.2-fold greater difference in methylation, whereas clams ‘primed’ in low pH (initial pH 7.4 and 7.0, secondary pH 7.4) showed very little methylation change (−0.4, and −0.5, respectively, **Figure 4C**). The most significantly overrepresented terms from GO enrichment were organic anion transport (GO:0015711; p=0.007) and phosphatidylinositol-5-phosphate binding (GO:0010314; p=0.007) (**Table S14**). The significantly enriched GO terms were related to the Biological Processes cell cycle and proliferation, cell-cell signaling, developmental processes, protein metabolism, RNA metabolism, stress response, transport, other biological processes, and other metabolic processes (**Table S14**).

**Figure 4.**
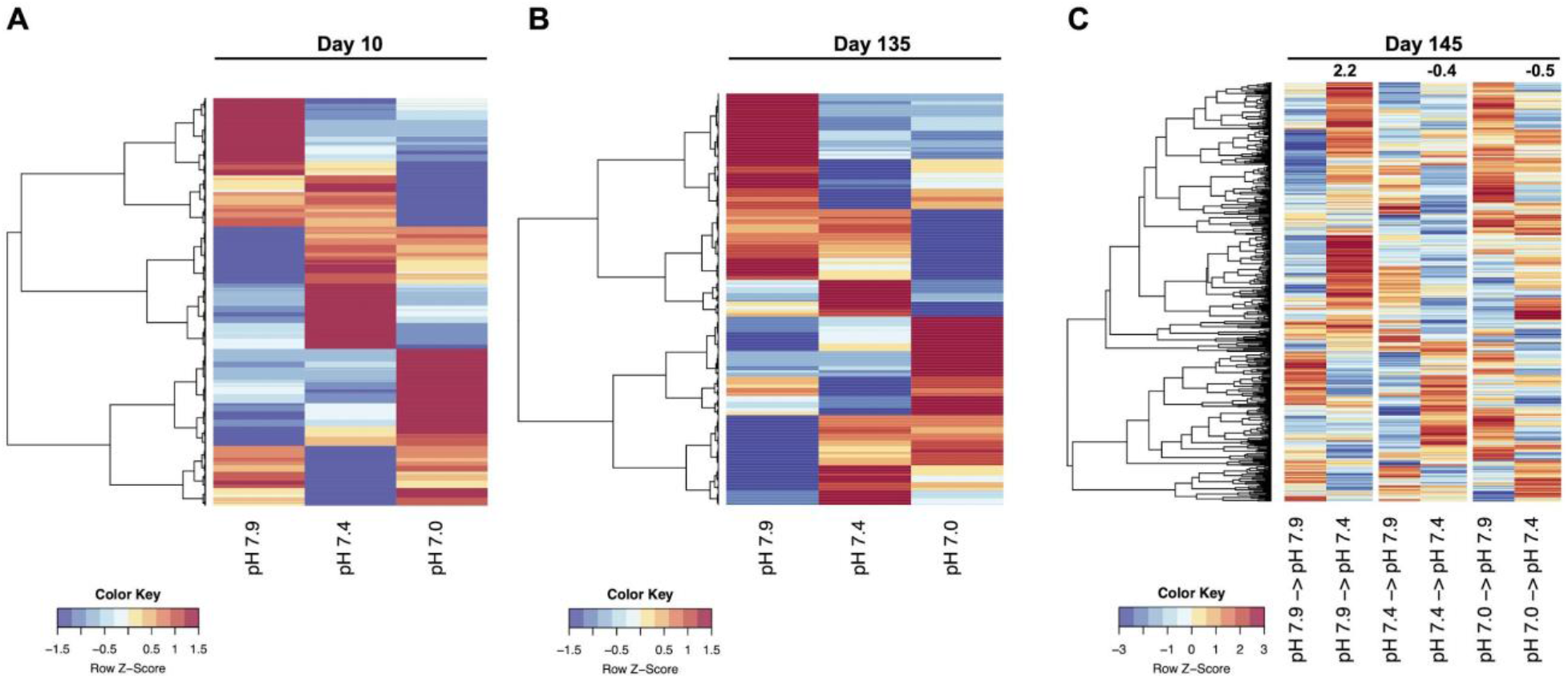
Differentially methylated genes after (**A**) initial pH exposure, (**B**) common garden, and (**C**) secondary pH exposure. Numbers above the paired columns in (**C**) represent the average change in methylation between the paired columns (separated by white space).

In examining persistent biological functions among DMGs from Day 10, Day 135, and Day 145, we found the Biological Processes cell cycle and proliferation, developmental processes, protein metabolism, stress response, transport, other biological processes, other metabolic processes persist across all three timepoints (**Figures S4** and **Figure 5**). Some Biological Processes only appear in a single, or pairs of time points (cell organization and biogenesis, cell-cell signaling, death, DNA metabolism, RNA metabolism, and signal transduction (**Figures S4**).

**Figure 5.**
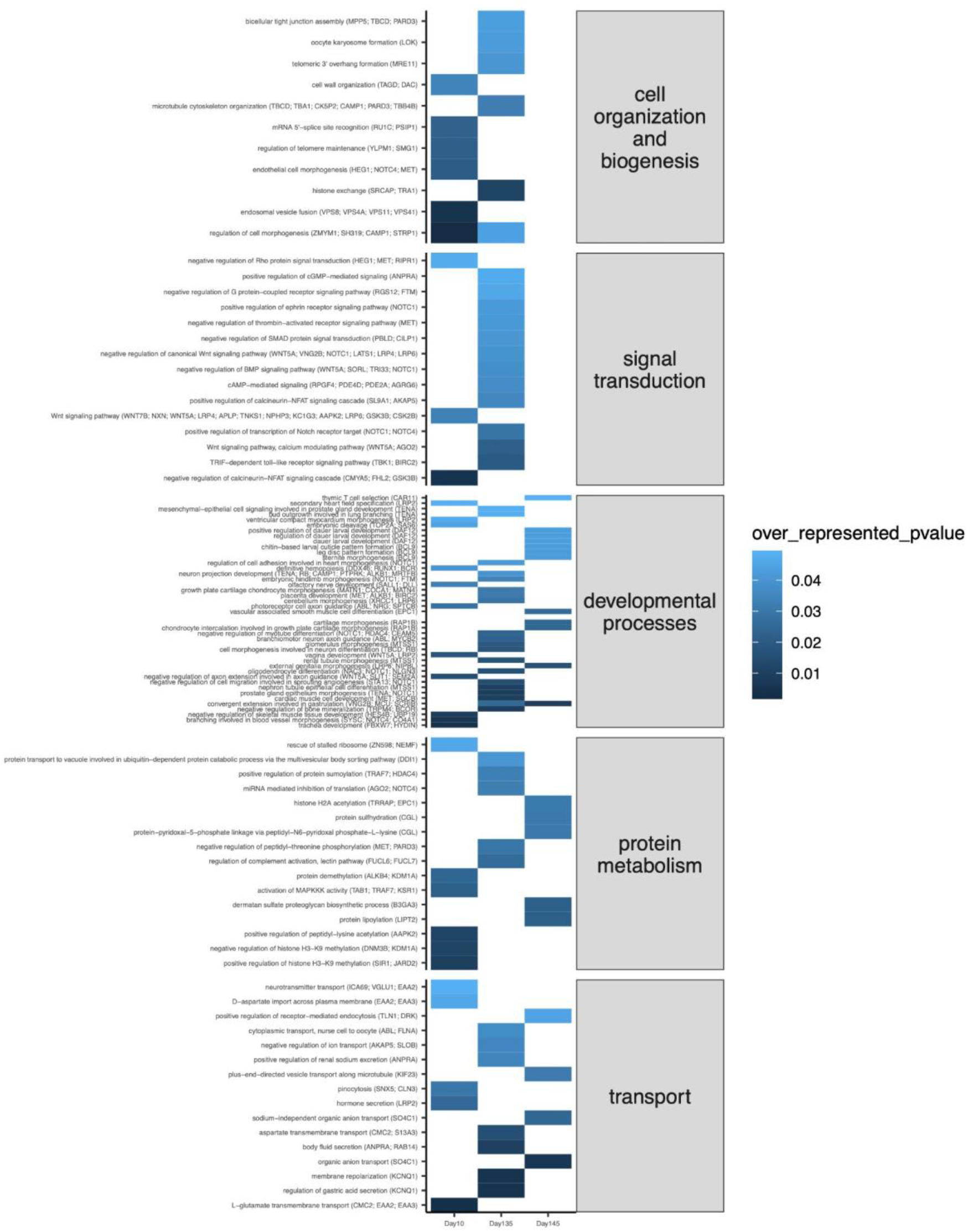
Subset of enriched GO biological processes for DMGs between treatment groups for Day10 (left column), Day 135 (middle column), and for the initial by secondary treatment interaction (acclimatization) for Day 145 (right column).

## DISCUSSION

### Overview of Findings

As anthropogenic climate change accelerates, both natural and aquaculture systems that humans depend on as protein resources are under threat. Understanding the mechanisms of tolerance and resilience of such fisheries taxa is therefore paramount to our capacity to support food security (Golden et al. 2016; Committee on World Food Security 2020). Here, we first generated a reference genome for *Panopea generosa* that provides an essential resource for the bivalve aquaculture field. Second, we conducted an experimental environmental hardening of clams at the juvenile hatchery stage, which resulted in a positive growth phenotype that lasted months after exposure, even in ambient conditions. Notably, when challenged again with low pH, those clams previously conditioned to low pH showed a buffered response (i.e., no change in relative size). On the basis of this phenotypic effect, we completed molecular characterization of gene regulatory epigenetic marks, specifically DNA methylation, through whole genome bisulfite sequencing, made possible by the generation of the reference genome. Our epigenetic analysis revealed differentially methylated genes (DMGs) following exposure and limited methylation change in the secondary exposure, if the clams were conditioned to low pH previously. Widespread differential methylation of genes in the Wnt signaling pathway, indicates a molecular basis for the phenotype in terms of cell fate and tissue formation, and a downstream cascade of differential methylation in cell organization and biogenesis, developmental processes, protein metabolism and transport. Collectively, these findings support an epigenetic component to compensatory growth in response to environmental hardening to low pH in juvenile geoduck clams.

### Genome and importance of genomic resources for aquaculture species

The generation of the Pacific geoduck draft genome as described here is critical to the mechanistic understanding of selective breeding, adaptation, acclimatization, and culture of these economically important and long lived clams. The draft genome assembly is on par with the limited number of genomes available for bivalves (Mun et al. 2017; Gómez-Chiarri et al. 2015; Zhang et al. 2012) in terms of size, gene number and GC content (Table 2). Specifically the *P. generaosa* genome size (942 Mb) is larger than the Pacific (Zhang et al. 2012) and Eastern oyster (Gómez-Chiarri et al. 2015), and more similar to *R. philippinarum* (1.23Gb; (Mun et al. 2017)). Both clams also have a larger number of putative chromosomes than the two oysters. In this *P. generaosa* assembly, approximately 80% of metazoan single copy orthologs identified via BUSCO analysis are accounted for in the 18 contigs. The *P. generaosa* genome can now be used as a resource and powerful tool for genetic marker discovery, gene isoform expression analysis, comparative genomics, shotgun proteomics (Timmins-Schiffman et al. 2020), and as a reference for epigenetic studies, such as the one described here.

### Location of DMRs and of methylation with respect to function

The variety of gene expression effects due to the location of methylation in the genome has yet to be fully elucidated (Roberts and Gavery 2012). For example, analysis of differentially methylated regions (DMRs) can provide insight into regional transcriptional capacity and epigenetic function because epigenetic function may result from combinatorial effects of neighboring CpGs, which often have highly correlated methylation levels (Zhang et al. 2015; Affinito et al. 2020; Korthauer et al. 2019). Additionally, DMRs are more frequently located near DEGs than individual differentially methylated CpGs and thus may be more biologically relevant (Aryee et al. 2014). Therefore we first explored DMRs here to determine if there was a higher order methylation architecture contributing to regional transcriptional regulation beyond that which has been documented at the gene level in invertebrates (i.e., correlation between gene body methylation and expression (Gavery and Roberts 2013), or between gene body methylation expression variability (Liew et al. 2018)). Introns on Day 145 held fewer DMRs and putative 3’ UTR regions on Day 10 held a greater number of DMRs than expected. Both intron and 3’ flanking regions can contain regulatory sequence, transcription factor binding sequence, and enhancer characteristics. With respect to gene expression regulation, intron methylation has also shown a positive correlation with expression in invertebrates (Anastasiadi et al. 2018) and may play a role in alternative splicing (Flores et al. 2012). This suggests methylation in these regions could play a regulatory role beyond the typical focus on gene body methylation in invertebrates (Dixon et al. 2018). Further experimentation is essential to identify the contribution of methylation in particular genomic features to expression regulation in invertebrates in general, and in geoduck clams specifically.

### Differentially methylated signal transduction genes indicate a role for the Wnt signal transduction pathways in impacting cytoskeletal, muscle, and shell/bone formation and organization

Given the annotation of many gene coding regions in DMRs and prior work identifying gene body methylation in invertebrates (Sarda et al. 2012; Roberts and Gavery 2012), we followed the regional analysis with a higher resolution gene level analysis to examine the methylation in protein coding genes and tested for enrichment in their functions. Differential methylation in genes is consistent with that observed in other marine invertebrates, particularly in Pacific oysters exposed to different temperatures (Wang et al. 2021), across development Pacific oysters (Riviere et al. 2017), and in Eastern oysters exposed to seawater conditions differing in pH (Venkataraman et al. 2020), where methylation differences have mainly been observed in exon and intron regions. In our study, DMGs were most abundant (1417 genes) after the initial 10 days of exposure, whereas after the ambient common garden period approximately half of that number showed differential methylation (797 genes). This suggests that there are “wash in, wash out” dynamics (Ptashne 2013; Burggren 2015) at the gene level, or that there may be a shift from DNA methylation as a primary regulatory feature to other mechanisms, such as histone modifications (Adrian-Kalchhauser et al. 2020; Li et al. 2018b).

The genes that were differentially methylated between pH treatments after 10 days into the primary exposure, and after the common garden (Day 135), resulted in the most widespread GO enrichment in signal transduction and cell organization, along with protein metabolism, transport, and developmental processes (**Figure 5**). Notably, the signal transduction enrichment included a variety of genes involved in the Wnt signaling pathway (**Figure 6**), with significant enrichment of seven different gene ontology terms related to Wnt signaling (**Table S10**). The Wnt signaling pathway is initiated by the binding of extracellular Wnt ligands to the cell surface receptor complex to initiate intracellular signal transduction (Buechling and Boutros 2011). This internal signal transduction cascade starts with the cytoplasmic phosphoprotein Dishevelled (Dsh) and from there can split into the three major pathways, the canonical Wnt β-catenin signaling pathway, or non-canonical pathways including the Planar Cell Polarity (PCP) and Wnt/Ca^2+^ pathways (Fig. 6). The canonical Wnt pathway is most well known for its role in body plan formation during development (Petersen and Reddien 2009), but also functions beyond development for cell proliferation of muscle, neuron, and osteoblast cells (Houschyar et al. 2018). In contrast, the two primary non-canonical Wnt pathways do not involve the activity of GSK3 on β-catenin and they function beyond early developmental stages to control cell morphogenesis and remodeling (Croce and McClay 2009).The planar cell polarity (PCP), as indicated by its name, is involved in cell polarity establishment and extension movement. In contrast, the Wnt/Ca^2+^ pathway regulates intracellular calcium levels to signal and regulate cytoskeletal dynamics, adhesion, and muscle growth (von Maltzahn et al. 2012). It is possible, however, for the same Wnt ligand to activate both the canonical and non-cannonical pathways (Kikuchi et al. 2009).

**Figure 6.**
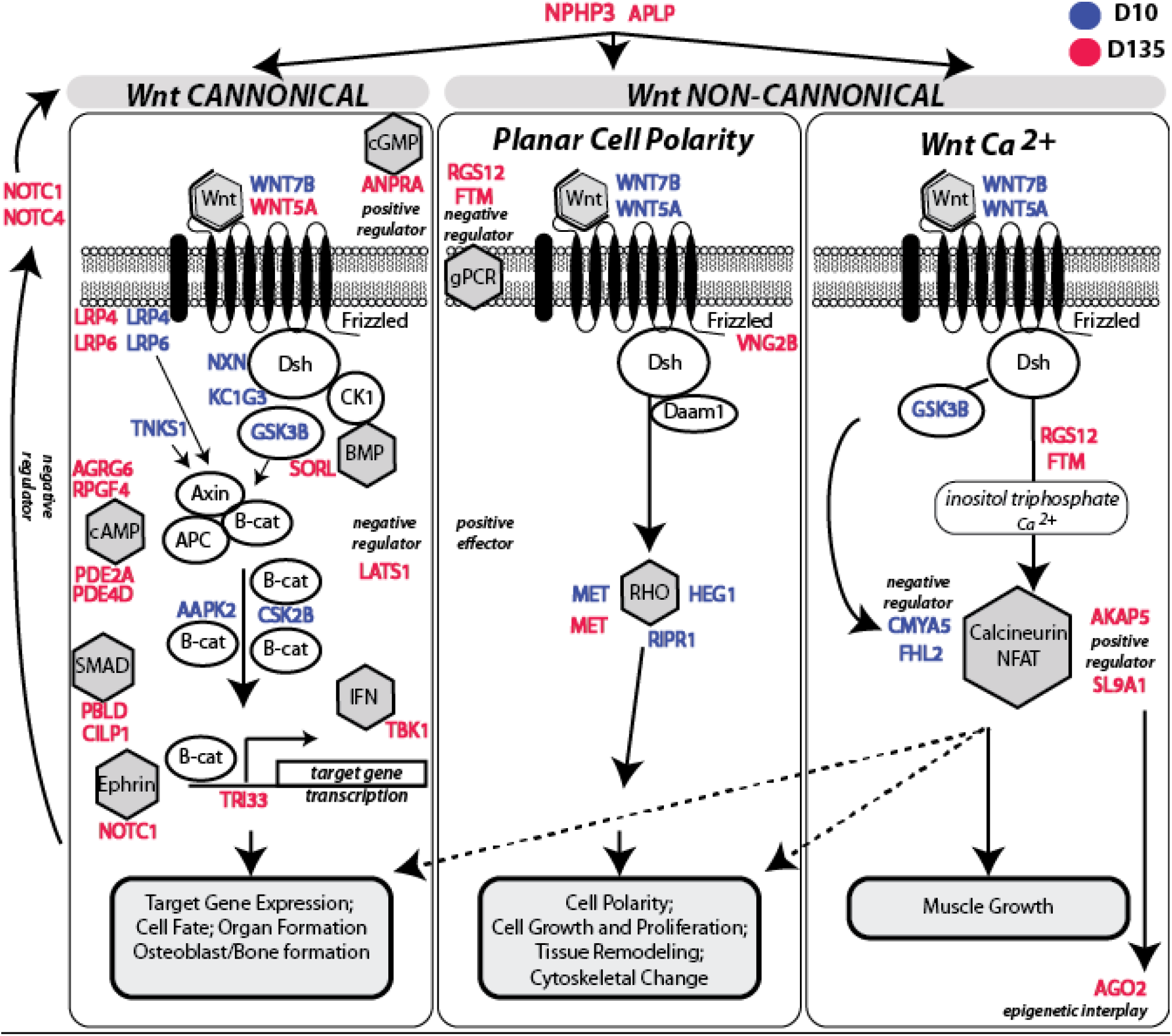
Differential signal transduction following exposure is linked to phenotype. Schematic of the three Wnt signaling pathways including Wnt specific genes, as well as their linkage to the other genes identified in the GO enrichment of signal transduction, leading to downstream consequences for cell and tissue formation, cell polarity and cytoskeletal change, and muscle growth. Genes that are shown in color indicate those that were differentially methylated and contributing to the significantly enriched GO term of signal transduction. Genes listed in blue indicate they contribute to significant GO term enrichment at Day 10 and genes listed red indicate they contribute to significant GO term enrichment at D135. Dsh=cytoplasmic phosphoprotein Dishevelled; Axin; APC=Adenomatous Polyposis Coli; CK1= Casein kinase 1; B-cat=β-catenin; Daam1= Formin protein. Gray octagons indicate signaling pathways. cGMP = cyclic guanosine monophosphate; cAMP = cyclic adenosine monophosphate; BMP = bone morphogenetic protein; SMAD = mothers against decapentaplegic homolog; IFN = interferon; EPH = ephrin; gPCR = G protein-coupled receptor; Rho= a GTPase; calcineurin-NFAT = calcineurin and nuclear factor of activated T cells.

In our study, there is evidence for a role of DNA methylation in all three Wnt pathways and therefore signal transduction (Fig. 6) linked to the clam growth phenotype. The presence of differential methylation in WNT5A and WNT7B along with coreceptors LRP4 and LRP6 indicates a role for the canonical signaling pathway, with ultimate consequences for cellular and skeletal structural processes (Fig. 6). This is supported by the presence of differential methylation of NXN (nucleoredoxin, a negative regulator of Wnt), which acts to provide a pool of inactive Dsh, the first step in the Wnt/β-catenin signaling pathway (Funato et al. 2010). Also KC1G3 (Casein kinase I isoform gamma-3) a serine/threonine kinase, is differentially methylated and binds and phosphorylates Dsh (Peters et al. 1999). Additionally TNKS1 (Poly [ADP-ribose] polymerase tankyrase-1) is differentially methylated and this gene further works to interact with AXIN (Axis inhibition protein)and form the β-catenin destruction complex with GSK3β. Further we see differential methylation in glycogen synthase kinase 3β (GSK3β), which has a role in phosphorylating β-catenin with subsequent impacts on signal propagation to transcription factors for target genes (Beurel et al. 2015). There was also differential methylation at Day 10 of AAPK2 (5’-AMP-activated protein kinase catalytic subunit alpha-2, an AMP-activated protein kinase, AMPK), which has a role for modulating β-catenin expression (Zhao et al. 2011), further supporting a pH treatment influence on the canonical Wnt pathway. Also indicative is the Day 10 differential methylation of CSK2β (Casein Kinase 2 Beta) between treatments.

The non-canonical Wnt PCP pathway transduces signals through Wnt ligand binding, which triggers Dsh, Daam1, and subsequently small Rho-like GTPases that enable regulation of the cytoskeleton (Fig. 6B). Differential methylation on Day 10 of MET (Hepatocyte growth factor receptor that activates Rho GTPases, (Royal et al. 2000)), HEG1 (Protein HEG homolog 1 a membrane protein that interacts with other proteins to inhibit Rho (de Kreuk et al. 2016)), and RIPR1 (Rho family-interacting cell polarization regulator 1 that acts to continue signal transduction and link Rho to additional kinases (Lv et al. 2022), indicates negative regulation of Rho protein signal transduction and provides further evidence for the role of PCP pathway and cytoskeletal effects in the clams following pH exposures.

Differential methylation in the Wnt signaling pathway is not only present after the acute exposure, but also after 135 days of ambient conditions in the common garden. In the canonical Wnt pathway, WNT7B, and the receptors LRP4 and LRP6 are differentially methylated. In the non-canonical pathways we also see DMGs on Day 135. In particular, differential methylation on Day 135 of Vang-like protein 2-B (VNG2B), a core PCP scaffolding protein (Hatakeyama et al. 2014) supports downstream PCP effects on cell adhesion and the cytoskeleton (Lindqvist et al. 2010). MET is likewise differentially methylated on Day 135. While there are the fewest DMGs in the Wnt/Ca^2+^ pathway on both Day 10 and Day 135, it remains possible there is a role for this in muscle or tissue growth. Gurr et al (Gurr et al. 2021) reported that following preconditioning of geoduck to low pH stress there was a subsequent increase in biomass. Specifically in our study, the support for the Wnt/Ca^2+^ pathway on Day 135 comes from differential methylation of RGS12 and FTM related to inositol triphophate Ca^2+^ linked to potential triggering of the calcineurin NFAT pathway where AKAP5 and SL9A1 are positive regulators.

The involvement of the Wnt signaling pathway is one way by which extracellular pH signals can be propagated to gene expression. Cellular acidosis has been shown in a variety of marine organisms exposed to ocean acidification (Kaniewska et al. 2012). Notably, H^+^-ATPase (V-ATPase) has been implicated in acidificiation of cells and connected to Wnt signaling (Cruciat et al. 2010). Specifically, intracellular acidification may enhance, or even be required for the phosphorylation of the LRP6 reception (Cruciat et al. 2010). Additionally, there is evidence that acid base status modulates specification of mesoderm in human tail bud precursor cells via β-catenin feedbacks (Oginuma et al. 2020) following intracellular pH changes. In a process analogous to clam cell differentiation and skeletal formation, osteogenic differentiation in mammalian cells also involves Wnt signaling. Specifically, when human bone marrow mesenchymal stem cells are induced to undergo osteogenic differentiation, DNA methylation increases. Within the set of genes with dynamics methylation, KEGG analysis identified enrichment of genes involved in the Wnt signaling cascade (Cao et al. 2018). Similar to the differential methylation seen after low pH exposures in Day 10 and Day 135, induction of osteogenic capacity in cells also results in changes in DNA methylation in Wnt and LRP-5/6 in the canonical Wnt/β-catenin pathway (Cao et al. 2018), suggesting a function of key players in the Wnt signaling pathway may influence osteogenesis and therefore calcification capacity. Indeed, genes involved in Wnt signaling have been suggested to have a role in shell formation in other bivalve species such as the pearl oyster (Gao et al. 2016). Taken together, the differential methylation genes involved in, or linked to Wnt signaling supports a linkage between DNA methylation in the canonical and non-cannonical pathways and the compensatory growth and buffering of shell size effects in the environmentally hardened clam phenotype.

### DNA methylation and Transcriptional Regulation in Invertebrates

While RNASeq was not conducted in our study, transcriptional regulation via DNA methylation has been described in other enriched gene ontologies of differential methylated features in Hong Kong oyster (*C. hongkongensis*) larvae exposed to low pH (Lim et al. 2021), as well as ontologies associated with protein modification and development. In Eastern oyster (*C. virginica*) gonad tissue, with differences in methylation found in several genes involved in protein ubiquitination and biomineralization (Venkataraman et al. 2020). In a complementary work on the same species (*C. virginica*), researchers looking at DNA methylation identified differential methylation in genes in mantle tissue DNA that are involved in biomineralization and stress response (Downey-Wall et al. 2020). However, when DNA methylation was characterized over time in *C. virginica*, the only functional overlap among genes containing differentially methylated loci were a suite of genes associated with the THO complex, a nuclear structure involved in transcription elongation and mRNA maturation (Downey-Wall et al. 2020). Taken together these data from bivalves demonstrate that low pH can influence methylation in genes associated with a variety of biological processes that are likely developmental and tissue specific. Collectively, there appears to be a consistency that alteration in DNA methylation in genes associated with transcriptional regulation is common upon exposure to ocean acidification conditions in bivalves.

Further work is needed here to identify the exact mode of action, signaling cascade, and epigenetic interplay (Adrian-Kalchhauser et al. 2020) involved in epigenetic “memory” in shellfish. In particular using long read sequencing technologies to sequence RNA and DNA from the same nucleic acid pools would provide data to test for alternative splicing and long read scaffold data providing a methylation signal without bisulfite sequencing (e.g., Nanopore, (Dimond et al. 2021)), or a combination of such approaches (Spektor et al. 2019).

### Conclusions and future needs

Genomic resources are essential for understanding mechanisms driving acclimatization and adaptation potential. Here we provide a key genomic resource for the geoduck clam, a species of high economic value. The value of this resource is clearly demonstrated here as a functional framework for the emerging field of environmental epigenetics. Future efforts could be focused on improving the completeness of the genome while also moving toward pan-genome and core-genome analysis (Whibley et al. 2021).

For the first time we have demonstrated an underlying alteration in the DNA methylation landscape that coincides with a beneficial phenotypic carry-over effect in a bivalve. These epigenetic findings may provide an exploitable role in environmental manipulation that would benefit aquaculture (Gavery and Roberts 2017). While arguably one would not need to know the underlying epigenetic mechanism conferring the beneficial phenotype (e.g., differential methylation of genes within the signal transduction cascade), this information could be leveraged to optimize environmental manipulation scenarios. Future research would benefit from a greater understanding of the interplay of epigenetic mechanisms involved in environmental memory, as DNA methylation is one of several processes including histone modification and regulatory RNAs (Adrian-Kalchhauser et al. 2020; Eirin-Lopez and Putnam 2019). Within long lived sessile benthic invertebrates, it will also be essential to see how life history overlaid onto genetic and epigenetic landscapes will influence ecological and evolutionary outcomes (Putnam 2021). Taken together, harnessing both hatchery and environmental settings will provide a rich testbed for both mechanistic understanding of epigenetics, as well as the use of conditioning practices to enhance the resistance and resilience of marine invertebrates.

## METHODS

### Genome Sequencing

Genomic DNA was isolated from adductor muscle and ctenidia tissues from a single adult geoduck. Tissue was pulverized under liquid nitrogen and DNA isolated using Magattract whole blood DNA extraction protocol (Qiagen). Extracted DNA was run on a pulsed-field gel electrophoresis to confirm the majority of the fragments were ≥100kb and used to prepare two 10X Genomics libraries sequenced in duplicate according to the manufacturer’s instructions with sequencing performed on an Illumina HiSeq4000 system. Raw sequence reads are available on the National Center for Biotechnology Information (NCBI) Sequence Read Archive (SRA) under the following accession numbers: SRX5775746-SRX5775753. Supernova (v2.0) was used to generate an assembly in the pseudohap format.

Chromatin conformation capture data was generated using a Phase Genomics (Seattle, WA) Proximo Hi-C Animal Kit, which is a commercially available version of the Hi-C protocol (Lieberman-Aiden et al. 2009). Following the manufacturer’s instructions, intact cells from two samples from the same geoduck mention above were crosslinked using a formaldehyde solution, digested using the Sau3AI restriction enzyme, and proximity ligated with biotinylated nucleotides to create chimeric molecules composed of fragments from different regions of the genome that were physically proximal in vivo, but not necessarily genomically proximal. Continuing with the manufacturer’s protocol, molecules were pulled down with streptavidin beads and processed into an Illumina-compatible sequencing library. Sequencing was performed on an Illumina HiSeq4000 (NCBI SRA: SRX1667267, SRX1667269, SRX1667275, SRX1667277, SRX1667279, SRX1667308, SRX5581791, SRX5775746 - SRX5775753).

Reads were aligned to the initial assembly of 10x Genomics reads following the manufacturer’s recommendations (Sullivan 2019). Briefly, reads were aligned using BWA-MEM (Li and Durbin 2010) with the −5SP and −t 8 options specified, and all other options default. SAMBLASTER(Faust and Hall 2014) was used to flag PCR duplicates, which were later excluded from analysis. Alignments were then filtered with samtools (Li et al. 2009) using the −F 2304 filtering flag to remove non-primary and secondary alignments.

Phase Genomics’ Proximo Hi-C genome scaffolding platform was used to create chromosome-scale scaffolds from the corrected assembly as described in Bickhart et al. (Bickhart et al. 2017). As in the LACHESIS method (Burton et al. 2013), this process computes a contact frequency matrix from the aligned Hi-C read pairs, normalized by the number of Sau3AI restriction sites (GATC) on each contig, and constructs scaffolds in such a way as to optimize expected contact frequency and other statistical patterns in Hi-C data. Approximately 400,000 separate Proximo runs were performed to optimize the number of scaffolds and scaffold construction in order to make the scaffolds as concordant with the observed Hi-C data as possible. Finally, Juicebox (Durand et al. 2016; Rao et al. 2014) was used to correct scaffolding errors (primarily due to a combination of very short contigs and haplotigs present in the assembly) as well as introduce one new break into contig 349, which appeared to be an assembly misjoin. Genome “completeness” was evaluated using BUSCO (v3.2.2; (Waterhouse et al. 2018; Simão et al. 2015) using the metazoa_odb9 dataset.

### Genome Annotation

A geoduck transcriptome assembly ((Roberts et al. 2020); available at https://osf.io/tb5kz/) was generated using RNAseq data from ctenidia, gonad, and heart tissues (NCBI BioProject PRJNA646071), as well as whole larval and juvenile animals (NCBI BioProjects: PRJNA566166, PRJNA529226) and used for RNA evidence data. StringTie (1.3.6; (Pertea et al. 2015, 2016) was used to identify splice variants from the same RNAseq data. TransDecoder (Grabherr et al. 2011) was used to predict proteins from the transcriptome assembly ((Roberts et al. 2020); available at https://osf.io/c6mub/). Genome annotation was performed using the Genome Sequence Annotation Server (GenSAS; Humann et al, 2019). All annotation files are available in the Center for Open Science (OSF;(Roberts et al. 2020)).

All tools described below were run on the GenSAS pipeline. Repeat identification and masking was performed with RepeatMasker (open-4.07; (Smit, AFA, Hubley, R & Green, P 2013-2015) using NCBI search engine, speed/sensitivity option −q, and invrep.ref database for species selection, along with RepeatModeler (1.0.11; (Smit and Hubley 2008). Three different sets of sequence alignments were run including 1) BLASTn (2.7.1; (Altschul et al. 1990; Camacho et al. 2009) against the NCBI RefSeq Invertebrate nucleotide database, 2) BLASTx against predicted proteins identified in our geoduck transcriptome (https://osf.io/c6mub/), and 3) DIAMOND (0.9.22; (Buchfink et al. 2015) protein alignment against the SwissProt database. Structural analysis and gene prediction used the EMBOSS *getorf* tool for open reading frame identification (Rice et al. 2000), tRNAscan-SE for identification of tRNA genes (2.0; (Chan and Lowe 2019), RNAmmer for prediction of rRNA (Lagesen et al. 2007), BRAKER (2.1; (Hoff et al. 2019, 2016; Stanke et al. 2008, 2006a; Lomsadze et al. 2014; Li et al. 2009; Barnett et al. 2011) using the StringTie BAM alignments, GeneMarkES (4.33; (Lomsadze et al. 2005; Ter-Hovhannisyan et al. 2008), as well as Augustus (3.3.1; (Stanke et al. 2008, 2006a, 2006b; Stanke and Waack 2003; Stanke 2003) and SNAP (2006; (Korf 2004). Augustus was run two times: once using our geoduck transcriptome assembly (Augustus-01) and once with the StringTie BAM alignments (Augustus-02). Both runs utilized the default species setting (*Drosophila melanogaster)*. SNAP utilized the fly snap_hmm library. Gene consensus was performed using EvidenceModeler with the following evidence weight assignments: Augustus-01 = 1, Augustus-02= 2, BRAKER = 1, GeneMarkES = 1, SNAP = 1, BLASTx = 5, DIAMOND = 5, BLASTn = 10. BUSCO (v3; (Simão et al. 2015; Waterhouse et al. 2018) using the metazoa_odb9 database was run on each of the resulting gene models produced by Augustus-01, Augustus-02, BRAKER, EvidenceModeler, GeneMarkES and SNAP, with 68.4%, 5.5%, 71.5%, 6.1%, 65.2%, and 3.3% complete BUSCOs, respectively. BRAKER gene predictions were selected as the official gene set and refinement was performed with the Program to Assemble Spliced Alignments (PASA; 2.3.3; (Haas et al. 2003), utilizing our assembled transcriptome. Functional annotations using the PASA-refined gene set were conducted with BLASTp (2.7.1; (Altschul et al. 1990; Camacho et al. 2009) and DIAMOND (0.9.22; (Buchfink et al. 2015) against SwissProt database. InterProScan (5.29-68.0; (Jones et al. 2014), Pfam ((Jones et al. 2014) and SignalP (4.1; (Petersen et al. 2011) analysis were also performed. After annotation was completed, BUSCO (v3; (Simão et al. 2015; Waterhouse et al. 2018) was run again on the final set of annotated genes.

### Experiment

Geoduck juveniles were exposed to a three-component experiment (**Figure 2**) designed to test for: 1) the effects of initial exposure to ocean acidification, 2) the potential for latent effects in an ambient common garden due to initial exposure, and 3) the potential for initial exposure to provide preconditioning and acclimatization to a secondary exposure. The experiments were conducted at the Kenneth K. Chew Center for Shellfish Research and Restoration in Manchester, WA. Juvenile geoduck clams were received at ~3 month of age (Taylor Hatchery Quilcene, WA) on 16 March 2016. Animals were placed in each of two 5L replicate tanks per treatment (35 × 21 × 12cm, lxwxh). The geoducks were allowed to bury themselves as desired in ~4cm of graded sand of the same source and grade from which the clams were collected at the hatchery. Temperature was monitored continuously using an Avtech system (TMP-DOT-SEN-AVTECH; accuracy = ± 2°C, **Figure S5 B**).

### Experimental Control System and Seawater Chemistry Analysis

The experiments were conducted using a flow-through, pH-stat system, with 1μm filtered seawater. The pH in the header tanks was continuously monitored with DuraFET pH probes (**Figure S5 A**) (Honeywell, Morristown NJ, USA) that fed pH and temperature data back into a solenoid controlled injection system. The pH set points were achieved by the injection of ambient air or pure CO_2_ into water lines of the header tanks, which continually cycled water from the bottom to the top of each header for even mixing and equilibration and delivery to the treatment tanks.

Seawater chemistry was assessed in each tank following best practices standards (Riebesell et al. 2011). Seawater pH, salinity and temperature were measured in each tank using handheld probes. pH was measured in mV (resolution = 0.01, DG115SC glass probe, Mettler Toledo, Ohio, USA) and calculated for the *in situ* temperature against a linear regression of a tris standard (Batch 2/14/16, salinity 27.5) as a function of temperature. Temperature was measured simultaneously, with a traceable digital thermometer (Accuracy: ±0.05°C, resolution: 0.001°, temperature range: –50 to 150°C, VWR, USA), and salinity with a traceable digital portable conductivity meter (Accuracy: 0.3%, Temperature Range: –30.0 to 130.0°C, VWR, USA). In combination with the probe measurements, 120 ml water samples were collected and stored in sealed borosilicate glass bottles and poisoned with 75μl of HgCl for total alkalinity analysis. Total alkalinity (TA) was measured using an open cell gran titration (Dickson SOP3; (Dickson et al. 2007) on a Mettler Toledo T50 automated titrator. While water samples for TA were collected throughout the experiment, the majority of the sample bottles suffered damage in storage. Therefore TA was calculated from the available measurements during the experiment and the mean value was applied to calculations of carbonate chemistry parameters throughout the experiments. Carbonate chemistry parameters (**Table S15**) were calculated using the seacarb package in R (Gattuso et al. 2016), within measured input parameters of pH (total scale), total alkalinity (μmol kg sw^−1^, salinity (psu), and temperature (°C), using constants of Kf from Perez and Fraga (Perez and Fraga 1987), Ks from Dickson (Dickson 1990) and K1 and K2 from Lueker et al (Lueker et al. 2000).

### Initial Exposure Conditions

For the initial exposure, treatments were set to pH 7.9 (ambient), pH 7.4, and pH 7.0. Treatment water was delivered through pressure compensating drippers to each tank at a rate of 9.6±0.1 LPH (mean±sem, n=6). Clams were fed a mix of diatoms and flagellates (*Cheatoceros sp*., *Cheatoceros muelleri*, *Pavlova pinguis, Tisochrysis lutea*) at a concentration of 65,123 ± 3,630 cells ml^−1^. To calculate shell size, samples were photographed lying flat with a size scale on days 0 (n=4 per treatment), 10 (n= 8 per treatment), and 23 (n= 8 per treatment). Shell size was assessed by measuring the length (longest distance parallel to the hinge), width (distance from hinge to the ventral edge, perpendicular to length), and area (planar surface area) of clams in photographs using ImageJ (Schneider et al. 2012). Samples for molecular analysis were snap frozen or stored in RNALater for use in molecular analysis and stored at −20 to −80°C until processing.

### Common Garden Conditions

At the end of the 23 days of exposure, the six tanks of clams were transferred to an ambient common garden condition. The geoducks were allowed to bury themselves as desired in ~4cm of graded sand of the same source and grade from which the clams were collected at the hatchery, at a density of 32 - 38 clams bin^−1^ (6 bins to track history). Ambient hatchery temperature (14-15°C) and pH (~7.9) were monitored continuously with an Avtech probe and Durafet pH sensor, respectively (Figure S1). After 28 days in the indoor common garden, clams were photographed for shell size analysis (n=16 per initial treatment group) as described above. Clams were pooled by initial treatment and transferred to three tanks (18 l) with mesh covered water exchange holes and placed hanging from a dock in Manchester, WA. Geoduck juveniles were held at ambient bay conditions (~14°C). After 84 days, clams were photographed for shell size analysis (n=28, 53, 54, for the initial treatments of pH 8.0, 7.4, and 7.0, respectively) and samples were collected for molecular analysis (n=8 per initial treatment group) by placing them in RNALater and freezing at −80°C.

### Second Exposure Conditions

Juvenile geoduck clams initially exposed to pH 7.9, 7.4, and 7.0 were returned to the hatchery for a second exposure to two levels of initial exposure, pH 7.9 and 7.4 (Figure 1). Animals were placed in replicate trays (~2.8L:15.7 × 15.7 × 12.1cm, lxwxh) per treatment. The geoducks were allowed to bury themselves as desired in ~4cm of the same graded sand used throughout the experiment. Sand was sterilized by autoclaving prior to use in indoor tanks. Treatment water was delivered through pressure compensating drippers to each tank at a rate of 3.14±0.09 (mean±sem, n=36) LPH. Clams were fed a mix of diatoms and flagellates (*Cheatoceros sp*., *Cheatoceros muelleri*, *Pavlova pinguis, Tisochrysis lutea*) at a concentration of 23,721±2,005 cells ml^−1^. Clams were measured and sampled after 10 and 23 days for both shell size and molecular analysis in each of the two secondary pH treatments, as described above.

### Shell Size Analysis

To measure clam shell size the length and width dimensions were quantified using ImageJ (Schneider et al. 2012). Relative shell size was determined by first calculating the group mean shell size then dividing each shell size measurement by its respective group mean. Linear modeling was used to test hypotheses 1 and 2, with the fixed factors of treatment and time, and for hypothesis 3 using the fixed factor of initial pH x secondary pH. Assumptions of normality and homoscedasticity were tested with graphical examination of the model residuals.

### DNA Methylation Library Preparation

For methylation characterization, DNA was extracted from 52 samples from 3 timepoints (Day 10, Day 135, Day 145) for n=4 clams for each treatment (Figure 2) using the Qiagen DNeasy Blood and Tissue Kit (Qiagen USA) according to manufacturer’s instructions with slight modifications. Briefly, samples were ground to a powder on liquid nitrogen and ~25mg of sample was incubated with lysis buffer (ATL) and proteinase K at 56°C for 1hr, mixing several times throughout. Samples were incubated an additional 10 minutes following addition of buffer AL. Prior to addition to the columns, 200μl of 100% ethanol was added to each sample and instructions were followed to complete the extraction. Samples were eluted in 125-200μl of buffer AE. Samples were quantified using Qubit BR dsDNA kit and quality checked using 1.5% TAE gel. Enzymatically sheared DNA was used to prepare libraries using the whole genome library preparation method. DNA samples were spiked with 0.5% (w/w) unmethylated lambda DNA (Promega cat: D1501) and incubated overnight at 37°C with MSPI (20U μl-1 NEB cat: R0106L) and 10x NEBuffer2 (NEB cat: B7002S) for digestion at the cut site of CC in the sequence CCGG. Digested samples were processed with the EZ DNA Methylation-Gold Kit (Zymo Research Catalog Nos. D5005) for bisulfite conversion according to manufacturer’s instructions with a 30μl input sample and a 12μl elution. Sample quality and fragment size was assessed using the Agilent 2011 Bioanalyzer RNA pico chip. Samples passing quality control were then used for Illumina library preparation Illumina TruSeq DNA Methylation library prep kit according to manufacturer’s instructions. Samples were prepared for multiplex sequencing using the Illumina DNA Methylation Kit Barcodes (Illumina Cat #: EGIDX81312). The resulting libraries were quality controlled using the Qubit dsDNA high sensitivity kit for quantification and the Agilent Bioanalyzer DNA High Sensitivity Kit. Libraries that passed QC (n=52) were pooled in equal molar concentrations and sequenced on the Illumina HiSeq 2500 (2×100bp) at GENEWIZ (NJ, USA).

### Sequence file processing and methylation calling

The fastq files (NCBI SRA: PRJNA566166) were checked for initial quality using FastQC (Andrews and Others 2010) with the paired option and filtered and trimmed using TrimGalore! (v0.6.4) (Krueger 2015) with the default settings, the `--paired` option, and clipping 8 bases from the 5’ and 3’ ends of both read 1 and read 2 following recommendations for Illumina TruSeq DNA Methylation Kit in the Bismark User Guide v0.22.2 (Roberts et al. 2020). To determine cytosine methylation, individual samples were mapped (to the *in silico* bisulfite converted geoduck genome described above) using Bismark (Krueger and Andrews 2011) with the option of `-score_min L,0,-0.6` (bowtie2 2.3.4.1; (Langmead and Salzberg 2012); samtools 1.9; (Li et al. 2009)). These same parameters were used to map the spiked lambda phage DNA to the lambda genome (Enterobacteria phage lambda complete genome Genbank Accession number J02459) to determine conversion efficiency in each sample. The resulting BAM files were deduplicated and methylation information extracted using Bismark. Deduplicated BAM files were also sorted and indexed using samtools to facilitate visualization. In addition, the `coverage2cytosine` function in Bismark was implemented using `--merge_CpG` and `--zero_based` to generate coverage files used to create bedgraph files and tab-delimited inputs for differentially methylated gene analysis.

### Differentially methylated region analysis

To identify differentially methylated genomic regions, the MethylPy (Schultz et al. 2015) workflow was used on tab-separated files containing all CpGs with 5x coverage to make four independent comparisons: all ambient samples, all samples from day 10, all samples from day 135, and all samples from day 145. First, the `DMRfind` function was used to call differentially methylated CpGs within each sample by testing each for a statistical difference from the estimated methylation variation due to genetics (Schultz et al. 2015). For merging differentially methylated CpGs into regions, the setting `--min-num-dms 3` was used to specify a minimum number of three differentially methylated CpGs required for a region and the setting `--dmr-max-dist 250` was used to require adjacent differentially methylated CpGs to be maximally 250 base pairs apart. The identified methylated regions were further filtered for coverage across at least 3 out of 4 samples (individuals) within a treatment group. For each of the four group comparisons, percent methylation data for methylated regions within each sample were arcsine square root transformed and a one-way ANOVA was run on each region to test for the effect of time (all ambient samples comparison) or of pH (Day 10, Day 135) on methylation by region. A 2-way ANOVA was used to test the effect of initial and secondary exposure (Day 145 sample comparisons) on region methylation. Regions showing differential methylation across groups were considered significantly differentially methylated if they had ANOVA *P* value less than 0.05. No FDR correction was applied to counter the loss of power from applying an ANOVA to percentage rather than count data. Heatmaps showing the percent methylation of significantly differentially methylated regions (DMRs) were generated using the `heatmap.2` function of the R package gplots (Gregory R. Warnes, Ben Bolker, Lodewijk Bonebakker, Robert Gentleman, Wolfgang Huber, Andy Liaw, Thomas Lumley, Martin Maechler, Arni Magnusson, Steffen Moeller, Marc Schwartz and Bill Venables 2020) using a Pearson correlation distance matrix and average clustering. To visualize size distributions of DMRs from each comparison, boxplots of DMR lengths were generated with `ggplot2`. To visualize DMR distribution across the genome, DMRs were plotted against scaffolds. To visualize DMR distribution across scaffold positions, relative scaffold position for each DMR was calculated by dividing the DMR start position by the length of the scaffold and DMRs were plotted against relative scaffold position. DMRs were next mapped to genomic features. Putative promoter (1000 base pairs upstream the transcription start site) and 3’ flanking region (2000 base pairs downstream the termination site) feature files were generated using the bedtools flank function. Next, feature files were concatenated into one master bed file and features were binned into 2000 base pair windows (the average maximum methylated region size across all comparisons) using the bedtools `windowmaker` function. Background regions were generated by first compiling all CpGs that satisfied the following requirements: 1) at least 5x coverage in each sample, 2) covered by at least three out of four samples per treatment group or time point, 3) covered by all groups in each comparison. Compiled CpGs for each comparison were then mapped to the binned genomic features using the bedtools `intersect` function. A 2000-base pair feature region was considered covered if it contained a minimum of three compiled CpGs. The reason for binning features and requiring three compiled CpGs within each 2000 base pair region was to generate background regions that were similar to the requirements for regions that DMRs were called from. DMRs were also mapped to the binned genomic features using the bedtools `intersect` function. The proportion of specific features with DMRs out of all features with DMRs was compared to the proportion of specific features in background regions out of all features in background regions using a chi square test of proportions. Feature proportions were considered significantly different if they had an FDR-corrected p.value of 0.1 or less. Stacked bar plots were generated with `ggplot2` in R.

### Differentially Methylated Gene Analysis

Differentially methylated gene analysis was also performed using the tab-delimited files generated from the `coverage2cytosine` function in Bismark filtered for CpG positions with a 5x coverage minimum. Bedtools `MultiIntersectBed` function (bedtools version C, (Quinlan and Hall 2010)) was used to identify positions sequenced in all samples, at all timepoints, for use in the statistical analysis of differentially methylated genes. Following filtering, all positions were joined with the gene feature track (Roberts et al. 2020) by matching the scaffold and position. The resulting dataset of all CpGs found at 5x coverage in all samples, in gene regions only, was further filtered to genes with at least 5 CpGs and this dataset was used for statistical analysis to test for differentially methylated genes (DMGs). To test the hypothesis that DNA methylation does not differ through time in ambient conditions, DNA methylation per gene under ambient conditions was modeled as a function of Time (fixed) for each gene. To test the hypothesis that DNA methylation does not differ between treatments after 10 days of exposure (Day 10), DNA methylation was modeled as a function of seawater pH (fixed, three levels). To test the hypothesis that DNA methylation does not differ between treatments after several months of ambient grow-out in a common garden (Day 135), DNA methylation was modeled as a function of initial seawater pH exposure (fixed, three levels). To test the hypothesis that initial exposure to low pH modulated DNA methylation in subsequent exposures (Day 145), DNA methylation was modeled as a function of initial seawater pH (fixed), secondary seawater pH (fixed). Heatmaps showing the percent methylation of significantly differentially methylated genes (DMGs) were generated using the `heatmap.2` function of the R package gplots (Gregory R. Warnes, Ben Bolker, Lodewijk Bonebakker, Robert Gentleman, Wolfgang Huber, Andy Liaw, Thomas Lumley, Martin Maechler, Arni Magnusson, Steffen Moeller, Marc Schwartz and Bill Venables 2020) using a Pearson correlation distance matrix and average clustering. Gene ontology (GO) enrichment analysis was completed for each set of DMGs due to treatment on Day10 and Day145, and for the set of DMGs from the interaction of Initial x Secondary treatments using the package goseq (Young et al 2010) in R (R Core Team 2017), using the background set of all genes identified for the DMG dataset as described above. A tab-delimited file mapping GO identifiers to 37 GO slim terms from Molecular Function (MF), Biological Process (BP), and Cellular Component (CC) categories was retrieved from the Mouse Genome Database (MGD) (Bult et al. 2019) at the Mouse Genome Informatics website, The Jackson Laboratory, Bar Harbor, Maine, World Wide Web (URL: http://www.informatics.jax.org/gotools/MGI_GO_Slim.html), July 2017.

## DATA ACCESS

The genomic DNA, transcriptomic, and DNA bisulfite sequence data generated in this study have been submitted to the NCBI BioProject database (https://www.ncbi.nlm.nih.gov/bioproject/) under accession numbers PRJNA316601; PRJNA529226 and PRJNA646071; and PRJNA566166, respectively. All intermediate data files and raw experimental data and statistical code are also available at OSF https://osf.io/yem8n/ (Roberts et al. 2020).

## ACKNOWLEDGMENTS

This work was funded in part through a grant from the Foundation for Food and Agriculture Research; Grant ID: 554012, Development of Environmental Conditioning Practices to Decrease Impacts of Climate Change on Shellfish Aquaculture. The content of this publication is solely the responsibility of the authors and does not necessarily represent the official views of the Foundation for Food and Agriculture Research.

