## supplemental material for "Dynamic DNA methylation contributes to carryover effects and beneficial acclimatization in geoduck clams"

**Supplemental Information Putnam Trigg et al Submission**

https://github.com/hputnam/Geoduck_Meth/tree/master/Supplemental

**Figure S1.** Size and genome location of differentially methylated regions.

**Figure S2.** Differentially methylated regions (DMRs) in the geoduck clam genome

**Figure S3.** Genomic features overlapping with DMRs and all regions in the genome

**Figure S4.** Enriched GO Biological Processes for DMGs from each sampling time point

**Figure S5.** Physical Conditions monitored in the tanks show (A) the experimental pH as measured by Durafet sensors and (B) Avtech temperature sensors within the experimental control system.

---

**Table S1.** Repeat content of the Pgenerosa_v1.0 genome, determined by RepeatMasker.

**Table S2.** Shell area 3-way ANOVA results

**Table S3.** Summary table of trimming statistics

**Table S4.** Summary table of alignment statistics

**Table S5.** Initial pH exposure effect DMR ANOVA results. Significant differentially methylated regions among pH treatments at Day 10

**Table S6.** Initial pH exposure effect after common garden DMR ANOVA results. Significant differentially methylated regions among the initial pH treatments following the ambient common garden (Day 135)

**Table S7.** Secondary pH exposure x time effect DMR ANOVA results. Significant differentially methylated regions among the initial and secondary pH treatments following the ambient common garden (Day 145)

**Table S8.** Chi square test results for proportion of DMRs in genomic features (Intergenic, exon, intron, putative 3' UTR, putative promoter, repeat region, and tRNA).

**Table S9.** Significant differentially methylated genes between the pH treatments at Day 10

**Table S10.** GO enrichment analysis of significant differentially methylated genes between between the pH treatments at Day 10

**Table S11.** Significant differentially methylated genes for the initial by secondary treatment interaction at Day 135 (following ambient common garden)

**Table S12.** GO enrichment analysis of significant differentially methylated genes for the initial by secondary treatment interaction at Day 135 (following ambient common garden)

**Table S13.** Significant differentially methylated genes for the initial by secondary treatment interaction at Day 145 (after 10 days of secondary $pCO_2$ exposure)

**Table S14.** GO enrichment analysis of significant differentially methylated genes for the initial by secondary treatment interaction at Day 145 (after 10 days of secondary $pCO_2$ exposure)

**Table S15.** Seawater chemistry results

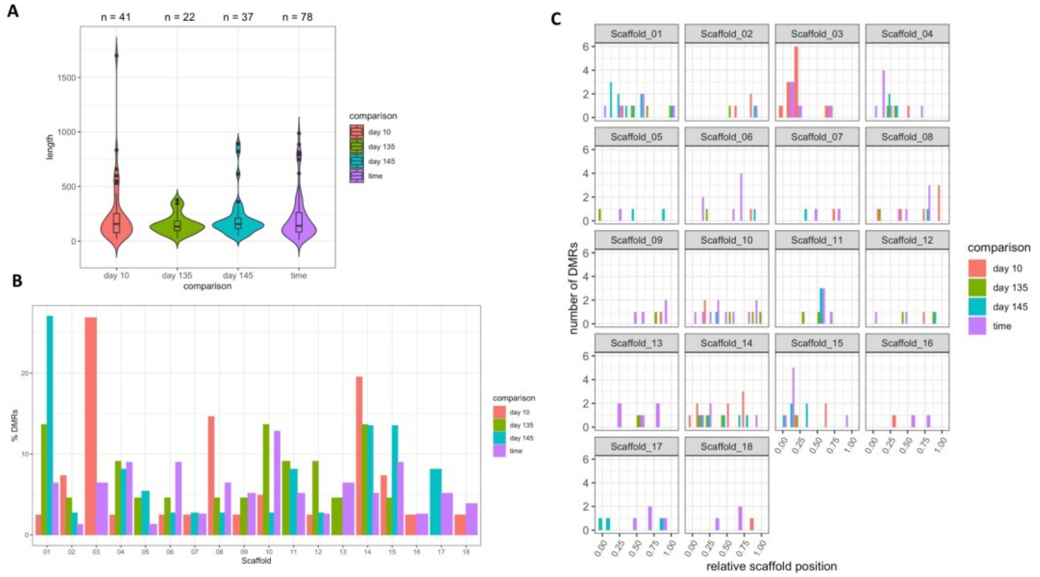


**Figure S1.** Size and genome location of DMRs. A) Violin plots showing the distribution of DMR lengths (in base pairs) across different sample comparisons (day 10, all samples from day 10; day 135, all samples from day 135; day 145, all samples from day 145; time, all ambient samples over time). Number of DMRs is listed across the top of the plot. B) Distribution of DMRs from different comparisons across genome scaffolds. Y axis is the proportion of DMRs in each scaffold out of the total number of DMRs in each comparison expressed as a percentage. C) Relative scaffold position of DMRs from different comparisons across genome scaffolds.

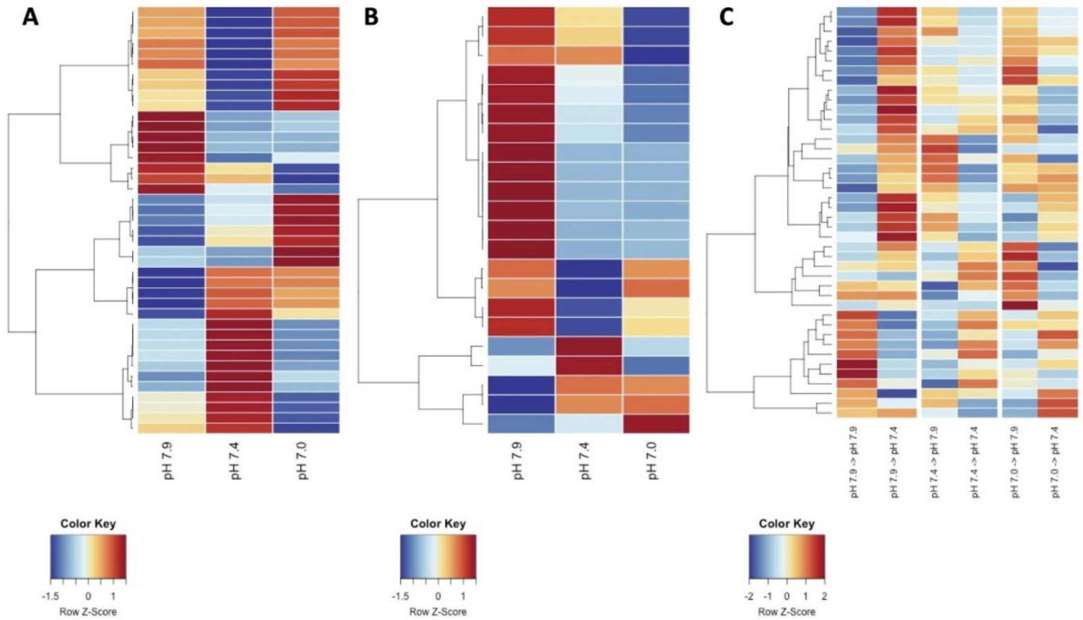


**Figure S2.** Differentially methylated regions (DMRs) in the geoduck clam genome after (**A**) 10 days of the initial high $pCO_2$ exposure (experimental day 10) (**B**) 112 days of common garden exposure (experimental day 135), and (**C**) 10 days of the secondary high $pCO_2$ exposure (experimental day 145). Heatmap color indicates the experimental group mean percent methylation of regions normalized by row (Z-Score).

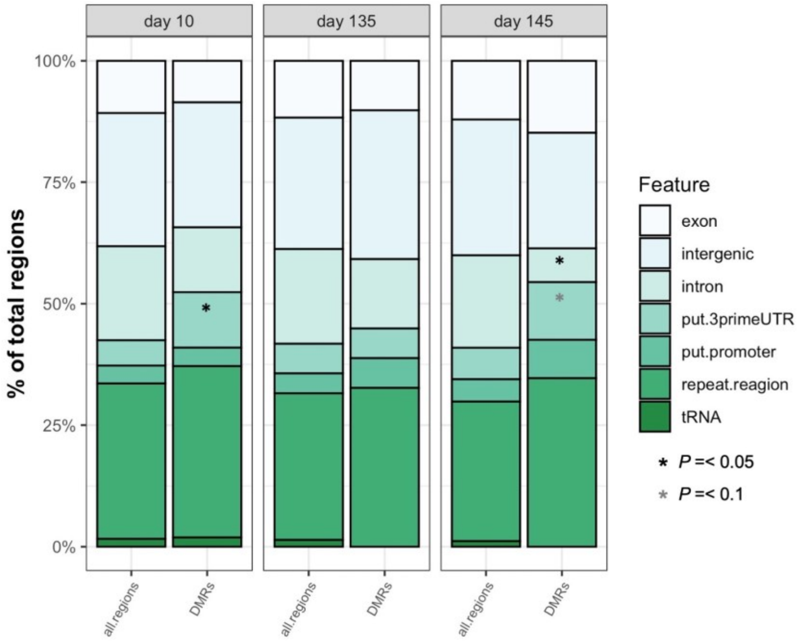


***Figure S3.*** *Genomic features overlapping with DMRs and all regions in the genome covered by samples from specific experimental timepoints. Asterisks indicate a significant FDR-corrected P value from a chi square test of proportions.*

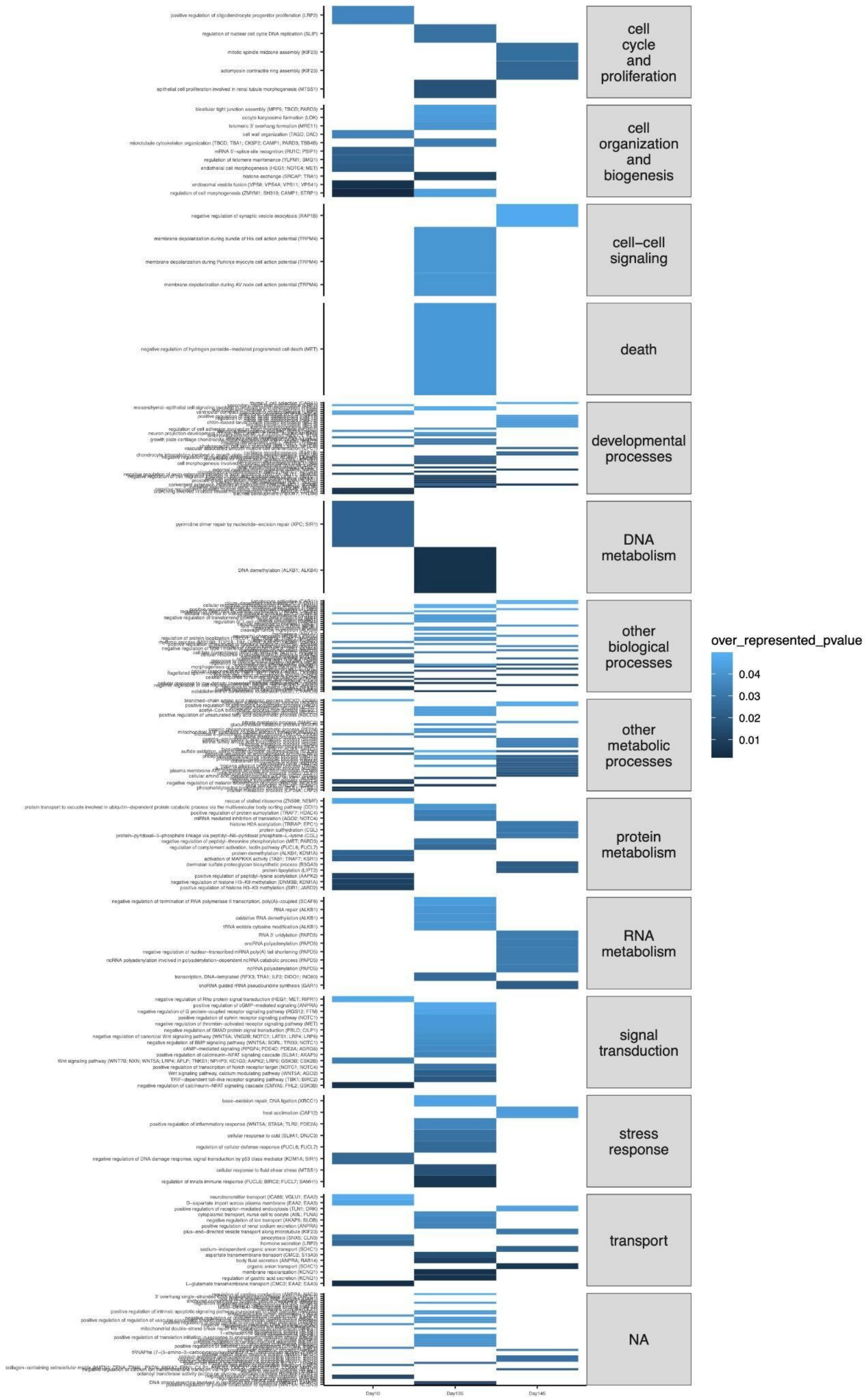


**Figure S4.** Enriched GO biological processes for DMGs between treatment groups for Day10 (left column), Day 135 (middle column), and for the initial by secondary treatment interaction (acclimatization) for Day 145 (right column).

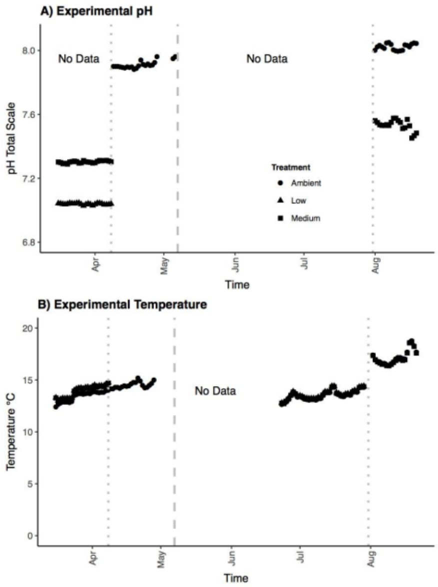


**Figure S5.** Physical Conditions monitored in the tanks show (A) the experimental pH as measured by Durafet sensors and (B) temperature within the experimental control system.
